# Synaptic plasticity deficits via aberrant engagement of metaplasticity in the hippocampus of PS19 mice

**DOI:** 10.64898/2026.06.30.734148

**Authors:** Shruthi Sateesh, Barbara J. Logan, Owen D. Jones, Wickliffe C. Abraham

**Author notes:** **Correspondence**: Dr Shruthi Sateesh, **Email**.

## Abstract

Tauopathy is characterized by progressive synaptic failure and neuroinflammation, yet the laminar-specific nature of these disruptions remains poorly understood. We investigated hippocampal functional integrity and glial reactivity in 8–10-month-old PS19 (P301S) mice. Electrophysiological recordings in the CA1 stratum radiatum revealed an unexpected increase in basal synaptic transmission despite profound deficits in both maintenance and early induction of the LTP phase. Conversely, the dentate gyrus exhibited reduced basal transmission and impaired LTP maintenance, alongside significant paired-pulse plasticity changes not observed in CA1. Furthermore, we demonstrate that transregional metaplasticity, as driven by prior activity in the stratum oriens (SO) in a way that inhibits subsequent LTP in wild-type mice, is occluded in PS19 mice. These data suggest that the tauopathic hippocampus exists in a “metaplastic” state, which inhibits future LTP. Immunofluorescence studies revealed that while astrogliosis and microglial activation were pan-hippocampal, specific neuroinflammatory markers exhibited striking laminar specificity. Mean fluorescence intensity for the neuroinflammatory astrocyte marker C3 was significantly upregulated only in the SO, and the lysosomal marker CD68 showed heightened occupancy specifically in the SO and stratum lacunosum-moleculare. Our findings indicate that tau pathology does not affect the hippocampus uniformly. Instead, it induces region-specific shifts in synaptic efficacy and a breakdown of metaplastic control that coincides with anatomically localized neuroinflammatory signaling.

## Introduction

In the healthy brain, long-term potentiation (LTP) and long-term depression (LTD) (Sateesh & Abraham, 2021) are dynamic processes governed in part by a history-dependent regulatory framework known as metaplasticity (Abraham & Bear, 1996). Described as the “plasticity of synaptic plasticity,” metaplasticity can serve a critical homeostatic function by shifting plasticity induction thresholds based on prior activity, to keep synaptic weights within a dynamic range and thereby help maintain network stability (Abraham & Bear, 1996). We have previously identified a form of heterodendritic metaplasticity in the hippocampus. Specifically, high-frequency “priming” stimulation of afferent inputs to the stratum oriens (SO) of area CA1 of the hippocampus triggers a synapse-specific inhibition of LTP and enhancement of LTD at apical dendritic synapses in the stratum radiatum (SR) (Hulme et al., 2012; Singh et al., 2022). This compartmentalized crosstalk, which also extends to the middle molecular layer (MML) of the dentate gyrus (DG) (Sateesh et al., 2026), appears to serve as a homeostatic brake to preserve the dynamic range of synaptic plasticity (Abraham, 2008; Hulme et al., 2013).

The spatial reach of this regulation, between hippocampal subregions and across the hippocampal fissure, suggests a non-neuronal, intercellular mechanism. Indeed, this form of metaplasticity occurs independently of postsynaptic membrane depolarization or the activation of ionotropic glutamatergic or GABAergic receptors, requiring instead a Ca^2+^ signal derived from an intracellular store (Hulme et al., 2012). This effect is blocked by gap junction inhibitors and peptide inhibitors of astrocytic connexin-43, suggesting a key role for the astrocyte network (Jones et al., 2013). Indeed, astrocytes respond directly to priming stimulation in the SO to subsequently inhibit LTP in the SR and MML (Hulme et al., 2013; Sateesh et al., 2026), supporting an astrocytic Ca^2+^-mediated signaling mechanism that regulates hippocampal LTP across broad spatiotemporal scales (Jones et al., 2023; Sateesh et al., 2026). Further, we have shown that a key signaling factor underpinning this form of metaplasticity is the gliotransmitter tumor necrosis factor (TNF) (Sateesh et al., 2026; Singh et al., 2019).

The hippocampus is acutely vulnerable in Alzheimer’s disease (AD) (Ebrahimi et al., 2025; Meftah & Gan, 2023). While classical views attribute cognitive decline primarily to late-stage synaptic loss and neuronal death, emerging evidence suggests that functional dysregulation—affecting both the immediate induction phase and the subsequent maintenance of LTP precedes overt neurodegeneration (Styr & Slutsky, 2018). Importantly, while metaplasticity mechanisms may help optimize encoding in the healthy hippocampus, in pathological states, they may become maladaptive, transforming from protective stabilizers into drivers of neural dysfunction (Abraham, 2008; Hulme et al., 2013).

In the APP/PS1 mouse model of amyloidosis, it is well established that metaplasticity mechanisms are aberrantly and chronically engaged (Jang & Chung, 2016; Megill et al., 2015; Singh et al., 2019). Of importance for this study, we have reported that, in these mice, there is a significant dampening of LTP in CA1 SR of aged APP/PS1 mice such that no further inhibition of LTP by SO priming can be elicited (Singh et al., 2019). Crucially, the LTP in these mice can be rescued by the bath-application of a TNF antibody, a treatment also known to block metaplasticity effects in normal animals (Singh et al., 2019). These data indicate that chronic neuroinflammation and dysregulated astrocyte-neuron interactions drive this aberrantly engaged metaplastic state that impairs LTP (Singh et al., 2019; Vincent et al., 2010).

While amyloid-beta (Aβ) is often viewed as the initial trigger for AD (Zhang et al., 2018), tau pathology, such as hyperphosphorylation and neurofibrillary tangles (NFTs), facilitates Aβ-induced impairment of LTP (Fá et al., 2016; Shipton et al., 2011). In addition, tau pathology correlates more strongly with synaptic plasticity impairments, cognitive decline, and clinical progression than the amyloid accumulation (Bejanin et al., 2017; Faldini et al., 2019; Iqbal et al., 2005). To date, the impact of pathological tau on the complex regulatory mechanisms of synaptic plasticity remains largely unexplored. To investigate this, we utilized the PS19 mouse model, which overexpresses the human P301S mutant tau (1N4R isoform). Unlike the common P301L mutation (0N4R), the P301S variant is characterized by a particularly low microtubule-affinity tau and aggressive aggregation kinetics, making it an ideal model for studying the transition from intracellular tau accumulation to neuroinflammatory cascades (Sahara & Yanai, 2023).

We hypothesized that the PS19 tauopathy model would exhibit an aberrantly engaged metaplasticity mechanism amid altered inflammatory signaling, mirroring observations in the APP/PS1 amyloid model (Di et al., 2016; Leyns & Holtzman, 2017; Rui et al., 2025; Vogels et al., 2019). In the present study, we demonstrate that PS19 mice exhibit significant LTP deficits both in SR of CA1 and in MML of DG, but with greater effects observed in CA1. These functional deficits were accompanied by alterations in input-output (I/O) curves and paired-pulse ratios (PPR), alongside significant reactive gliosis marked by increased immunolabelling for astrocytic glial fibrillary acidic protein (GFAP), the microglial marker Ionized calcium-binding adapter molecule 1 (IBA1), and the neuroinflammatory marker complement component 3 (C3). Similar to the APP/PS1 mice (Singh et al., 2019), the ability to metaplastically inhibit subsequent LTP was lost in both SR of CA1 and MML of DG. Our findings suggest that the hippocampus of P301S tau-overexpressing mice, similar to APP/PS1 mice, exhibits glial activation accompanied by aberrantly engaged metaplasticity in the hippocampal circuitry that impairs LTP prior to gross neurodegeneration.

## Materials and Methods

Experiments were conducted using 8–10-month-old male and female PS19 (P301S) transgenic mice (B6.Cg-Tg(Prnp-MAPT*P301S) PS19Vle/J; Strain #024841; The Jackson Laboratory, Bar Harbor, USA). Hemizygous male mice were bred with wild-type (WT) C57BL/6 female mice to generate transgenic offspring and their corresponding WT littermate controls. All animals were reared in specific-pathogen-free (SPF) conditions at the University of Otago Breeding Station. Following delivery to the research facility, mice were group-housed (2–3 per cage) in individually ventilated cages (IVC) in a controlled environment (23 ± 2°C; 12:12 h light/dark cycle, lights on at 6:00). Health status was monitored and recorded twice weekly using standardized monitoring sheets. All experimental objectives and animal handling procedures were approved by the University of Otago Animal Ethics Committee (AUP22-43) and performed in strict accordance with New Zealand animal welfare legislation. The study design and reporting adhere to the ARRIVE guidelines. Tissue samples were outsourced to Transnetyx (TN, USA) for genotyping.

Aged WT and PS19 transgenic mice were anesthetized with isoflurane and decapitated. Brains were rapidly dissected and submerged in ice-cold sucrose dissection solution (in mM: 210 sucrose, 26 NaHCO_3_, 2.5 KCl, 1.25 NaH_2_PO_4_, 0.5 CaCl_2_, 3 MgCl_2_, 20 D-glucose, saturated with 95% O_2_/5% CO_2_). Horizontal slices containing the middle hippocampus (350 μm) were prepared using a Leica VT 1200 S vibratome. Slices were incubated at the interface of air and artificial cerebrospinal fluid (ACSF, in mM: 124 NaCl, 3.2 KCl, 1.25 NaH_2_PO_4_, 26 NaHCO_3_, 2.5 CaCl_2_, 1.3 MgCl_2_, 10 D-glucose) at 32°C for 30 min, followed by room temperature until use. Post-incubation slices were submerged in a recording chamber perfused with ACSF at 2.5 ml/min and a temperature of 32.5°C ± 0.2 °C.

Prior to LTP experiments, IO curves and paired-pulse data were taken from SR and MML of each slice to assess basal synaptic transmission and paired-pulse plasticity, respectively. The average of three IO recordings (interpulse interval 30 s) was taken from each of six stimulus input increments (10, 20, 50, 100, 150, 200 µA). Paired-pulse tests were conducted using two pulses at varying inter-pulse intervals (20, 30, 50, 75, 100, 150, 200 ms). The average of three recordings, spaced by 30 s, at each interval was taken. The paired-pulse tests were taken at an input stimulation strength not sufficient to induce a population spike, to ensure that feedback inhibitory mechanisms were not engaged.

Field excitatory postsynaptic potentials (fEPSPs) were recorded from the SR of CA1 and the MML of the DG in the hippocampus, using previously described methods (Singh et al., 2022). Briefly, custom-built programmable stimulators evoked fEPSPs by delivering constant current diphasic pulses (100 μs per half-wave) via 50 μm Teflon-insulated tungsten monopolar wire stimulation electrodes. Pulled micropipette recording electrodes, filled with 2 M NaCl (1.5–2.5 MΩ), were placed 400 μm away from the stimulating electrode to record fEPSPs. For standard LTP experiments in the absence of priming, electrodes were placed in the SO (S1) and the test pathway (S2) within either the SR or MML; alternatively, two slices were recorded simultaneously in the test pathway. For priming experiments, sets of stimulating (S1 and S2) and recording electrodes (R1 and R2) were placed in both SO and SR (Fig. 1A), or SO and MML (Fig. 2A), with the stimulating electrodes and recording electrodes placed parallel to the relevant cell body layer. The criteria for including a slice in the 0065xperiment were a fEPSP response of at least 1.5 mV from SR in CA1 and MML in DG, and at least 0.5 mV in SO, to a 100 µA input. Baseline stimulation was then set at 40% of the fEPSP slope elicited at a stimulation strength of 200 µA.

**Figure 1:**
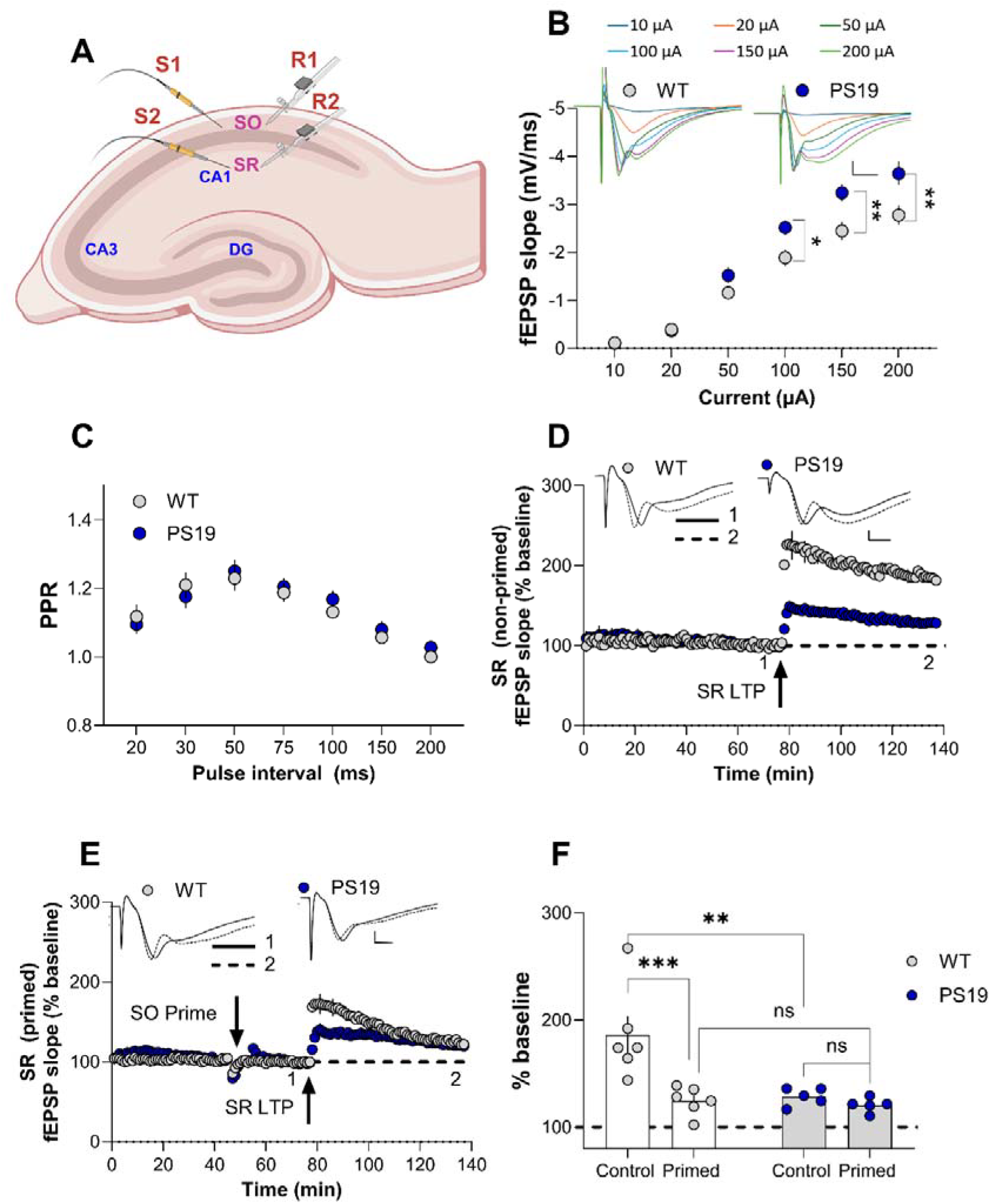
Impaired LTP and aberrant heterodendritic metaplasticity in CA1 SR of the PS19 mouse hippocampus. **(A)** Schematic of electrode placement in a mouse hippocampal slice, illustrating stimulation (S) and recording sites (R) in the SO (S1,R1) and SR (S2,R2). **(B)** Input-output (I/O) curves at SR synapses in PS19 and WT littermates across a range of stimulation intensities (10–200 µA). **(C)** Presynaptic short-term plasticity (paired-pulse facilitation) in SR was expressed as the mean ratio across inter-pulse intervals (20–200 ms). **(D,F)** LTP induced in SR under control conditions (in absence of SO priming). PS19 mice exhibited significant deficits in the induction and maintenance of LTP compared to WT mice. **(E,F)** Heterodendritic metaplasticity. SR LTP 30 min post-priming in SO was significantly reduced in WT mice, while no such reduction was observed in PS19 mice, for which LTP was already impaired, indicating aberrantly engaged metaplasticity. **(F)** Summary of SR LTP magnitude at the end of the experiment in the presence and absence of SO priming. Insets: Representative fEPSP waveforms recorded prior to induction (1) and at the conclusion of the experiment (2). Arrows indicate the timing of SO priming and SR LTP induction. Scale bars: **(B)** 1 mV, 5 ms; **(D,E)** 0.5 mV, 2 ms. All data presented as mean ± SEM; ns, p > 0.05; * p < 0.05; ** p < 0.01; *** p < 0.001.

**Figure 2:**
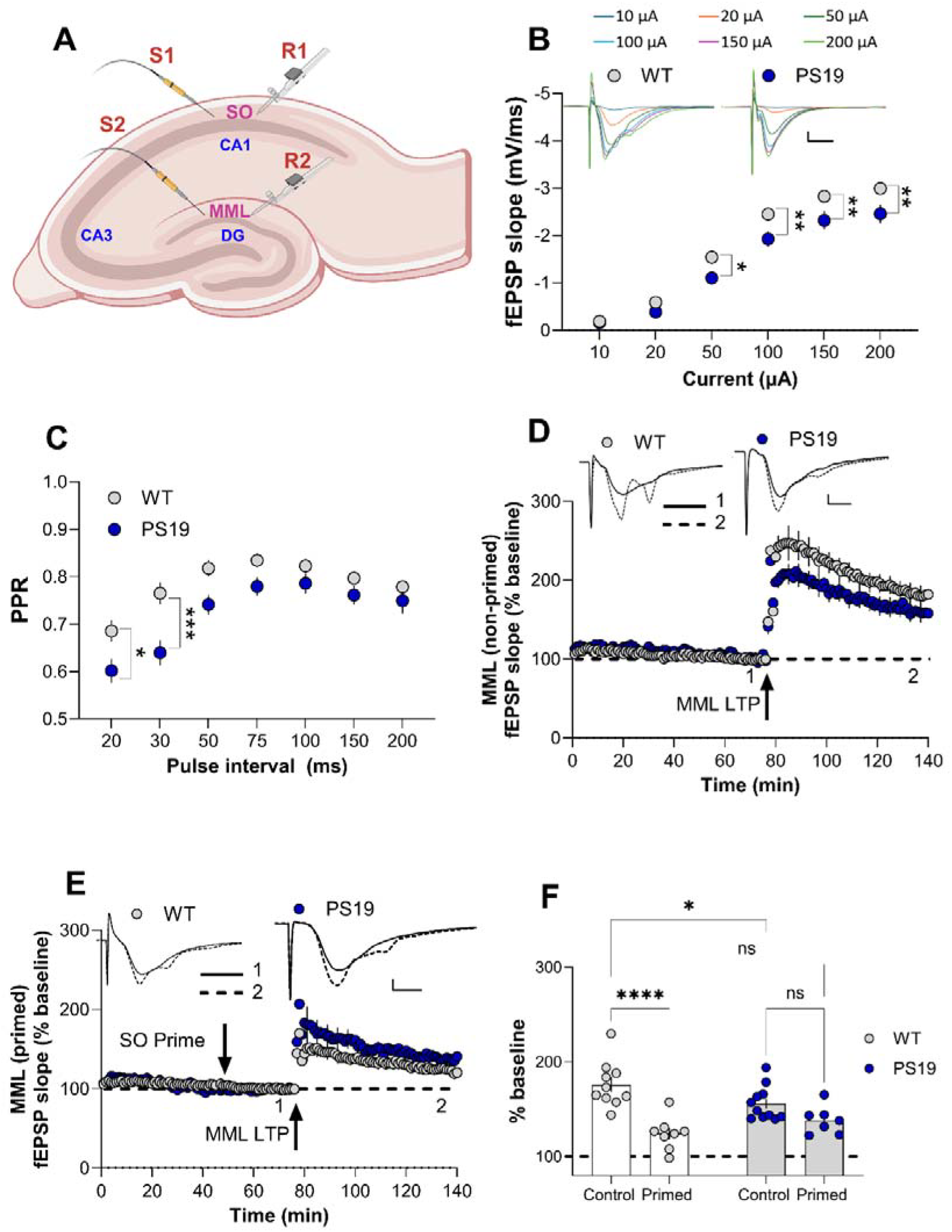
Impaired LTP and aberrant transregional metaplasticity in the DG MML of PS19 mouse hippocampus. **(A)** Schematic of electrode placement in an acute transverse mouse hippocampal slice, illustrating stimulation (S) and recording sites (R) in the SO (S1,R1) and MML (S2,R2). **(B)** Input-output (I/O) curves at MML synapses in PS19 and WT littermates across a range of stimulation intensities (10–200 µA). **(C)** Presynaptic short-term plasticity (paired-pulse depression) in MML was expressed as the mean ratio across inter-pulse intervals (20–200 ms). **(D,F)** LTP induced in MML under control conditions (in absence of SO priming). PS19 mice exhibited significant deficits in the maintenance of LTP compared to WT. **(E,F)** Transregional metaplasticity. MML LTP 30 min post-priming in SO was significantly reduced in WT mice, while no such reduction was observed in PS19 mice, for which LTP was already impaired, indicating aberrantly engaged metaplasticity. **(F)** Summary of MML LTP magnitude in the presence and absence of SO priming. Insets: Representative fEPSP waveforms recorded prior to induction (1) and at the conclusion of the experiment (2). Arrows indicate the timing of SO priming and MML LTP induction. Scale bars: **(B)**1 mV, 5 ms; **(D,E)** 0.5 mV, 2 ms. All data presented as mean ± SEM; ns, p > 0.05; * p < 0.05; ** p < 0.01; *** p < 0.001.

For SO priming experiments, baseline stimulation was delivered every 30 s, alternating between the priming stimulating electrode placed in SO and the test stimulating electrode in either SR or MML. Control experiments included 75-80 min of baseline stimulation followed by LTP induction either in SR or MML, and 60 min post-tetanization recording. SO priming stimulation, when given, occurred 30 min before conditioning stimulus in the test pathway and comprised of two trains of theta-burst stimulation (TBS) with 20 s between stimulation, with each train consisting of 10 bursts of 5 pulses at 100 Hz, 200 ms between bursts, at baseline stimulation amplitude; in SR, the same stimulation was repeated twice with a 30 s intertrain interval. LTP induction in DG MML used 4 trains of TBS comprising 10 bursts of 10 pulses at 100 Hz, 200 μs pulse duration, 200 ms interburst intervals, 30 s intertrain intervals in the presence of 0.2 μM gabazine to reduce GABA_A_ receptor-mediated inhibition.

### Immunofluorescence and image quantification

Different animals were used for the immunofluorescence studies. Following transcardial perfusion with a choline chloride-based solution (in mM: 92 choline chloride, 2.5 KCl, 12 NaH_2_PO_4_, 30 NaHCO_3_, 20 HEPES, 25 glucose, 2 L-ascorbic acid, 2 thiourea, 3 sodium pyruvate, 10 MgSO_4_.7H_2_O, 0.5 CaCl_2_.2H_2_O, 5 N-acetyl-L-cysteine), brains were harvested and post-fixed in 4% paraformaldehyde (4°C, 24 h), and subsequently cryoprotected in 30% sucrose (0.1 M PB containing 0.1% azide). Horizontal sections (40 µm) were collected using a cryostat (Leica Biosystems) and stored in ethylene glycol-based cryoprotectant at 4°C.

For immunolabeling, 3 sections/mouse, six sections apart, containing the mid-dorsal hippocampus were picked. The sections were washed in 0.1 M phosphate buffer (PB) to remove cryoprotectant, followed by antigen retrieval in 10 mM sodium citrate buffer (pH 6.0, 80°C, 1 h). Sections were blocked (3% Normal Goat Serum, 0.1% Triton X-100 in 0.1 M PB) and incubated overnight at 4°C with primary antibodies against GFAP (chicken anti GFAP: 1:4000, Invitrogen, PA1-10004 ), C3 (rat anti-C3: 1:500, Abcam, ab11862), Iba1 (rabbit anti-IBA1: 1:1000, Invitrogen, MA5-41239 ), and CD68 (rat anti CD68: 1:500, Invitrogen, MA5-16674). After washing, sections were incubated with Alexa Fluor-conjugated secondary antibodies: goat anti-Chicken, Alexa Fluor 555 (1:2000, Invitrogen, A32932), goat anti-rabbit, Alexa Fluor 555 (1:1000, Invitrogen, A21429), and goat-anti-rat, Alexa Fluor 647 (1:1000, Invitrogen, A48265) for 2 h at room temperature, then mounted using a custom antifade medium. Imaging was performed on a Nikon C2+ confocal microscope (20x objective, NA 0.75) with 2x optical zoom, focusing on the hippocampus. Images were deconvolved (NIS-Elements) and analyzed using ImageJ (v1.54q). Regions of interest (ROIs) were defined as the CA1 strata - SO, stratum pyramidale (SP), SR, and stratum lacunosum-moleculare (SLM), and the molecular layer (ML) and granule cell layers (GCL) in the DG **(Fig. 3C & 4C)**. Following 8-bit conversion and thresholding standardized to WT controls, the percentage area and mean intensity of immunopositive particles (> 4 µm^2^) were quantified and averaged across three sections for each animal.

**Figure 3:**
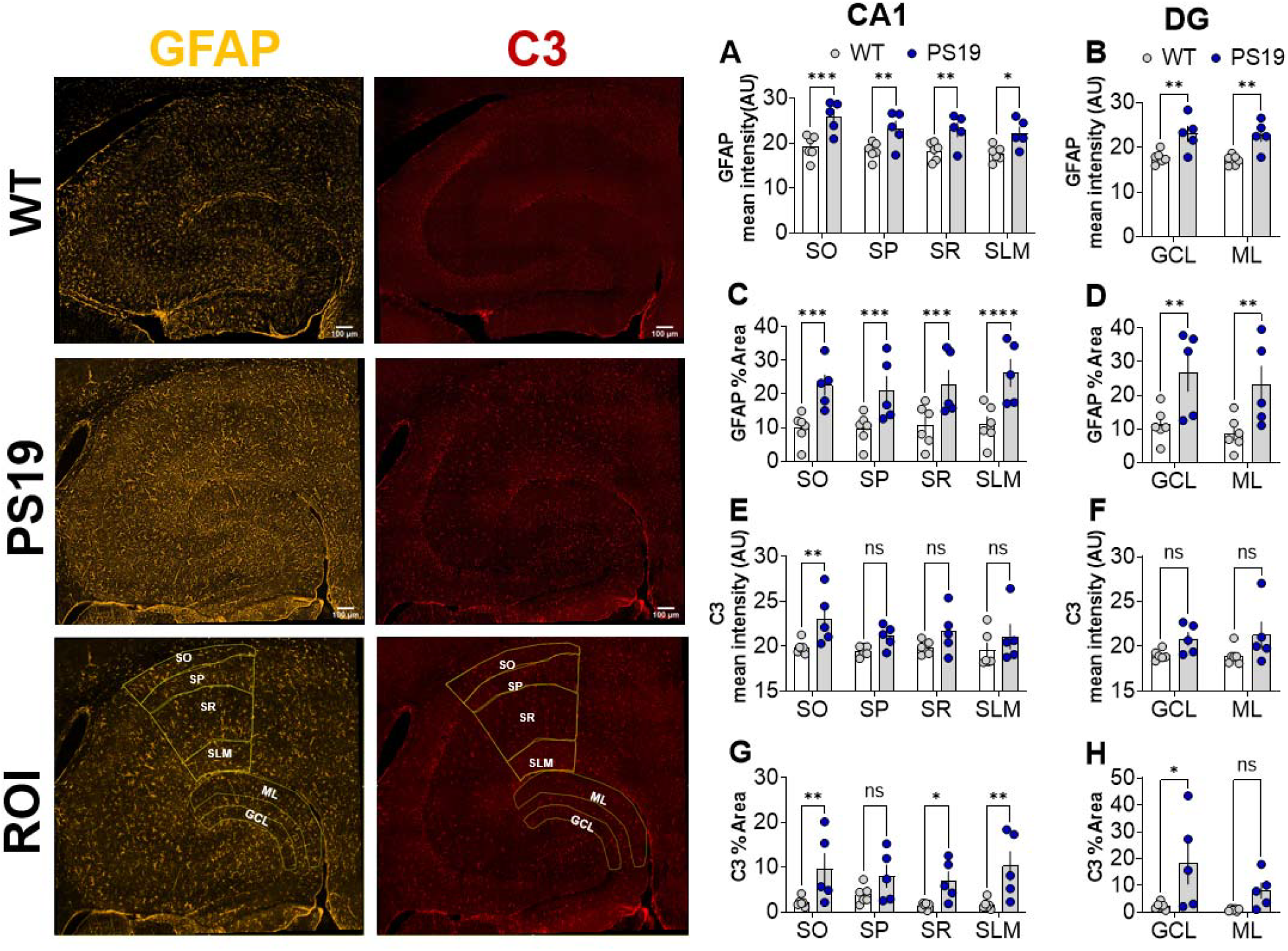
Astrocyte (GFAP) activation and C3 complement expression in the PS19 hippocampus. **(A-C)** Representative confocal images of GFAP expression (yellow) in the hippocampus of WT and PS19 mice. **(D-F)** Corresponding images showing C3 expression (red) across hippocampal subfields. **(C,F)** Regions of interest (ROI) used for laminar analysis, including the stratum oriens (SO), stratum pyramidale (SP), stratum radiatum (SR), and stratum lacunosum-moleculare (SLM) of the CA1, as well as the molecular layer (ML) and granule cell layer (GCL) of the dentate gyrus (DG). **(G-J)** Quantification of GFAP mean intensity and percentage area coverage reveals a significant increase in astrogliosis across all CA1 and DG layers in PS19 mice compared to WT littermates. **(K-N)** Quantification of C3 mean intensity and percentage area. PS19 mice exhibited elevated C3 levels, most notably in the SO, SR, and SLM of CA1 and the GCL of the DG, consistent with a reactive astrocyte phenotype. Scale bars: 100 µm. Individual data points are an average of three sections per animal (n = 5-6 mice per group). All data presented as mean ± SEM; ns, p > 0.05; * p < 0.05; ** p < 0.01; *** p < 0.001; **** p < 0.0001.

### Data analysis

Quantitative assessments and electrophysiological recordings were performed blinded to the genotypes. Recorded fEPSPs were analyzed offline using custom-built software (LabVIEW-based). The initial slopes of fEPSPs were measured, and the baseline value was defined as the average slope during the 10 min preceding LTP induction. All subsequent synaptic responses were expressed as a percentage of this baseline, with LTP magnitude calculated by averaging responses during the final 10 min of the experiment. The data for each group were pooled across animals, with individual slices being the experimental unit of analysis (n’s). Group sizes were *n* = 5-11, as based on our previous studies. As an a priori criterion, slices that exhibited changes in baseline responses⍰>⍰10% were excluded from analysis.

For immunofluorescence, the percentage area covered and thresholded mean intensity for GFAP, C3, Iba1, and CD68 were quantified by averaging three sections per mouse. Data were analyzed using GraphPad Prism (v10.6.1; San Diego, CA, USA). Statistical significance was determined using an unpaired Student’s t-test or two-way ANOVA for comparison between the genotypes, followed by Bonferroni’s, Fisher’s LSD, or Tukey’s post-hoc tests for multiple comparisons. The alpha level was set at 0.05. and all data are presented as mean ± SEM.

## Results

### PS19 mice have altered basal synaptic transmission in the CA1

To characterize the impact of tau overexpression on hippocampal functional integrity, we first assessed basal synaptic transmission at Schaffer collateral (SC)-commissural synapses within the SR of 8–10-month-old PS19 mice and WT littermates (**Fig. 1A**). An input-output analysis revealed significant main effects of genotype (F_(1, 120)_ = 22.1, *p* < 0.0001, **Fig. 1B**) along with a significant genotype x stimulus intensity interaction (F_(5, 120)_ = 3.0, *p* = 0.0143, **Fig. 1B**). Bonferroni’s post-hoc analyses demonstrated that PS19 mice exhibited significantly steeper fEPSP slopes at the higher stimulus intensities, specifically at 100 µA (*p* = 0.038), 150 µA (*p* = 0.004) and 200 µA (*p* = 0.0013) relative to WT littermates.

To determine whether alterations in presynaptic release probability correlated with these changes, we measured the PPR across various inter-pulse intervals (**Fig. 1C**). While a two-way ANOVA confirmed a significant effect of interval (F_(6, 91)_ = 10.3, *p* < 0.0001, **Fig. 1C**), there was no significant difference between genotypes (F_(1, 42)_ = 2.2, *p* = 0.15) and no genotype x interval interaction (F_(6, 42)_ = 0.71, *p* = 0.65, **Fig. 1C**).

### PS19 mice exhibit deficits in SR LTP and heterosynaptic metaplasticity

We next assessed LTP in CA1. We first examined the immediate response within the first 5 minutes post-TBS (early induction phase LTP) in the SR and found it to be significantly reduced in PS19 mice (WT: 220.7 ± 25.6%, *n* = 6; PS19: 139.9 ± 9.4%, *n* = 5, t_*(9)*_ =2.7, *p* = 0.02, **Fig.1D**). This deficit in the early induction phase was mirrored by a significant impairment in SR-LTP maintenance 1 hr later in PS19 mice compared to WT controls (**Fig. 1D,F**). A two-way ANOVA revealed a significant main effect of genotype (F_(1, 10)_ = 9.93, *p* = 0.01, **Fig. 1D,F**). Post-hoc analysis (Uncorrected Fisher’s LSD) confirmed that the magnitude of SR LTP was significantly reduced in PS19 mice (PS19: 128.6 ± 3.7%, *n* = 5, **Fig. 1D,F**) relative to WT controls (WT: 186.1 ± 17.5%, *n* = 6, *p* = 0.003, **Fig. 1D,F**). This substantial reduction in SR LTP was consistent with previous reports (Yoshiyama et al., 2007), indicating that tauopathy at 8–10 months severely disrupts the mechanisms required for hippocampal synaptic strengthening.

To investigate whether this impairment could be the result of aberrant engagement of the metaplasticity effect, 2xTBS priming stimulation was delivered to the SO pathway (S1), while conditioning stimulation was applied to the SR pathway (S2) 30 min later (**Fig. 1A**). Postsynaptic responses were monitored via recording electrodes R1 (SO) and R2 (SR). Consistent with our non-primed findings above, we observed a significant reduction in the early induction phase in the PS19 primed slices (PS19: 126.9 ± 7.1%, *n* = 5) compared to WT primed (WT: 166.9 ± 11.2%, *n* = 6, *t*_*(9)*_ =2.9, *p* = 0.01, **Fig.1E**). When measured 1 h later in WT mice, this SO-priming protocol induced robust heterodendritic inhibition of SR LTP (WT: 124.8 ± 5.4%, *n* = 6, **Fig. 1E,F**) compared to non-primed controls (*p* = 0.0003), replicating previously reported metaplasticity effects in CA1 SR of mice (Hulme et al., 2012). In contrast, PS19 mice failed to show further inhibition of SR LTP following SO priming (PS19: 120.7 ± 3%, *n* = 5, **Fig. 1E,F**). The magnitude of SR LTP in primed PS19 slices was not significantly different from non-primed PS19 slices (*p* = 0.64) or even compared to primed WT slices (*p* = 0.78). These data suggest an aberrant engagement of metaplasticity in the PS19 mice, occluding any further inhibitory effects of heterosynaptic priming.

We also assessed the degree of LTP in the SO priming pathway (S1). While the 2xTBS priming stimulus in SO induced robust LTP in WT mice (WT: 153.8 ± 14.32, *n* = 6, Fig. S1), LTP in the PS19 mice was significantly reduced (PS19: 111.14 ± 7.2%, *n* = 4, *t*_*(8)*_ =2.6, *p* = 0.03, Fig. S1), suggesting a widespread synaptic dysfunction in CA1 of the hippocampus in this mouse model. Note, however, that induction of LTP in SO is not a requirement for heterodendritic metaplasticity in CA1 (Hulme et al., 2012; Jones et al., 2013; Sateesh et al., 2026) and so the lesser LTP in PS19 mice does not account for the lack of a priming effect on LTP in SR.

### Presynaptic Deficits but spared LTP at MPP-DG Synapses

While synaptic dysfunction in the DG has been characterized in P301L and other tau-overexpressing models (Ash et al., 2025; Boekhoorn et al., 2006; Liu et al., 2017), the electrophysiological profile in the DG of aged PS19 (P301S) mice remains less defined. Here, we assessed synaptic efficacy by stimulating MPP inputs and recording fEPSPs in the MML of DG (**Fig. 2A**). Analysis of basal synaptic transmission revealed a significant reduction in fEPSP slopes in PS19 mice compared to WT controls (F_(1, 126)_ = 36.47, *p* < 0.0001, **Fig. 2B**), but there was no significant genotype X intensity interaction (F_(5,120)_ = 2.0, *p* = 0.07). Thus, in contrast to the SR of CA1, PS19 mice exhibited significantly reduced fEPSP slopes across various input current intensities: 50 µA (*p* = 0.03), 100 µA (*p* = 0.004), 150 µA (*p* = 0.006), and 200 µA (*p* = 0.003) compared to WT littermates.

To determine if alterations in presynaptic release probability contributed to these changes, and to confirm specific stimulation of the MPP, we measured the PPR across multiple inter-pulse intervals. In the DG, MPP stimulation is characteristically identified by paired-pulse depression (PPD). A two-way ANOVA confirmed a significant effect of interval (F_(6, 168)_ = 12.3, *p* < 0.0001, **Fig. 2C**) and a significant main effect of genotype (F_(1, 112)_ = 30.9, *p* < 0.0001), with a smaller PPR for the PS19 mice. Post-hoc analysis indicated that the PPR deficits in PS19 mice were most prominent at short inter-pulse intervals of 20 ms (*p* = 0.04, **Fig. 2C**) and 30 ms (*p* = 0.0003, **Fig. 2C**).

### PS19 mice exhibit deficits in MML LTP and trans-regional metaplasticity

To assess if synaptic plasticity was impaired in the DG of aged PS19 mice, we recorded LTP at DG-MML synapses. Unlike in the CA1 SR, the DG showed no significant difference in the early-induction phase of LTP between PS19 and WT mice (PS19: 191.7 ± 9.9%, *n* = 11; WT: 212.8 ± 8.9%, *n* = 10, *t*_*(19)*_=1.6, *p* = 0.13, **Fig. 2D**), suggesting that the initial induction mechanisms in the DG remain relatively resilient. Despite this preserved induction, a two-way ANOVA revealed a significant genotype X priming interaction for the maintenance of LTP (F_(1,32)_ = 6.9, *p* = 0.013). Post-hoc analysis showed that while TBS induced robust LTP in both groups, the magnitude of MML LTP maintenance was significantly attenuated in PS19 slices (155.9 ± 5.4%, *n* = 11, **Fig. 2D,F**) compared to the WT littermate controls (176.1 ± 7.6%, *n* = 10, *p* = 0.009; **Fig. 2D,F**), albeit to a lesser extent than in CA1.

We next investigated whether the transregional metaplasticity mechanisms were aberrantly engaged in the DG. Thus, priming stimulation was delivered to the CA1 SO (S1), followed by LTP induction in the DG MML 30 minutes later (S2; **Fig. 2A**). Consistent with our non-primed DG findings, we found no significant difference in the LTP induction phase between primed PS19 and WT slices (PS19: 178.6 ± 20.9%, *n* = 7; WT: 148.1 ± 16.1%, *n* = 7, *t*_*(12)*_ =1.2, *p* = 0.3, **Fig. 2E**).

When measured 1 h after SO priming, the MML LTP in WT mice was robustly inhibited (WT: 124.2 ± 6.1%, *n* = 7, **Fig. 2E,F**) compared to non-primed controls (*p* =0.03), confirming the transregional metaplasticity effect previously reported (Sateesh et al., 2026). In contrast, SO priming failed to further inhibit MML LTP in PS19 mice (PS19: 138.3 ± 5.6%, *n* = 7, **Fig. 2E,F**). The magnitude of MML LTP in primed PS19 slices was not significantly different from non-primed PS19 slices (*p* = 0.15) or even compared to primed WT slices (*p* =0.13). These results indicate that the metaplasticity mechanism is occluded in the aged PS19, likely due to its chronic engagement across the hippocampal fissure.

In the SO priming pathway, 2xTBS induced a homosynaptic LTP in PS19 mice (137.1 ± 16.4%, *n* =7, Fig. S2) that was no different from WT mice (136 ± 23.6%, *n* =7, *t*_*(12)*_ = 0.02, *p* = 0.98, Fig. S2).

### Laminar-Specific Glial Activation and Neuroinflammatory Profile in CA1

Previous findings from our group established that the metaplasticity is TNF-mediated and pathway-specific within the hippocampus, with synapses in the SR and MML exhibiting differential sensitivity to TNF priming compared to SO and SLM (Sateesh et al., 2026; Singh et al., 2022). Building on this, we investigated whether glial reactivity follows a similar laminar organization in the PS19 tauopathy model. Confocal immunofluorescence analysis revealed a robust state of astrogliosis across the CA1 and DG subfields, characterized by widespread increases in GFAP mean intensity and percent area coverage (**Fig. 3A-C, G-J**). A two-way ANOVA confirmed a significant main effect of genotype on GFAP mean intensity in CA1 (F_(1, 36)_ = 38.01, p <0.0001, **Fig. 3G**) and in the DG (F_(1, 8)_ = 32.5, *p* = 0.0005, **Fig. 3H**). Post-hoc tests indicated significant astrogliosis across all CA1 laminae, with robust intensity increases in the SO (WT: 19.2 ± 1.1, *n* = 6; PS19: 25.9 ± 1.5, *n* =5, *p* = 0.0003, **Fig. 3G**), SP (WT: 18.3 ± 0.8, *n* = 6; PS19: 23.2 ± 1.7, *n* = 5, *p* = 0.006, **Fig. 3G**), SR (WT: 18.2 ± 0.82, *n* = 6; PS19: 22.9 ± 1.6, *n* = 5, *p* = 0.008, **Fig. 3G**) and SLM (WT: 17.7 ± 0.7, *n* = 6; PS19: 22.2 ± 1.4, *n* = 5, *p* = 0.01, **Fig. 3G**). This was mirrored in the DG, where both the GCL (WT: 17.8 ± 0.6, *n* = 6; PS19: 23.0 ± 1.7, *n* = 5, *p* = 0.004, **Fig. 3H**) and the ML (WT: 17.2 ± 0.5, *n* = 6; PS19: 22.7 ± 1.5, *n* = 5, *p* = 0.003, **Fig. 3H**) showed significant elevations.

The upregulation of the intensity of labelled pixels was accompanied by a roughly doubling of labelled astrocyte occupancy (GFAP percentage area, **Fig. 3A-C, I-J**), with a significant main effect of genotype in both CA1 (F_(1, 16)_ = 90.30, *p* < 0.0001, **Fig. 3I**) and DG (F_(1, 8)_ = 32.52, *p* = 0.0005, **Fig. 3J**). The expansion was most pronounced in the SLM (WT: 11.0 ± 2.3%, *n* = 6; PS19: 26.3 ± 4.1%, *n* = 5, *p* < 0.0001, **Fig. 3I**), suggesting a high degree of astrocytic hypertrophy in the layer receiving distal inputs to the CA1 pyramidal cells. Robust increases were also noted in the SO (WT: 9.9 ± 1.8%, *n* = 6; PS19: 22.7 ± 3.1%, *n* = 5; *p* = 0.008, **Fig. 3I**), SR (WT: 10.7 ± 2.5%, *n* = 6; PS19: 22.9 ± 4.3%, *n* = 5, *p* = 0.0004, **Fig. 3I**) and SP (WT: 9.8 ± 1.9%, *n* = 6; PS19: 21.1 ± 4.2%, *n* = 5, *p* = 0.0006, **Fig. 3I**). Within the DG, both the GCL (WT: 11.3 ± 2%, *n* = 6; PS19: 26.9 ± 5.6%, *n* = 5, *p* = 0.004, **Fig. 3J**) and ML (WT: 8.7 ± 2%, *n* = 6; PS19: 23.1 ± 5.7%, *n* = 5, *p* = 0.006, **Fig. 3J**) exhibited significant increases in astrocytic percentage area coverage, suggesting that glial response in the PS19s represents a pan-hippocampal transformation of the astrocytic network.

In contrast, we found that expression of the neuroinflammatory astrocyte marker C3 displayed a more restricted profile (**Fig. 3D-F, K,L**). While genotype effects were significant in CA1 (F_(1, 16)_ = 14.1, *p* = 0.0017, **Fig. 3K**) as well as in the DG (F_(1, 18)_ = 6.5, *p* = 0.02, **Fig. 3L**), post-hoc analysis revealed that C3 protein upregulation was confined exclusively to the **SO** (WT: 19.9 ± 0.31, *n* = 6; PS19: 23.1 ± 1.3, *n* = 5, *p* = 0.009, **Fig. 3K**). No significant alterations in intensity were detected in the SP (*p* = 0.16, **Fig. 3K**), SR (*p* = 0.11, **Fig. 3K**), or SLM (*p* = 0.19, **Fig. 3K**) CA1 subfields or in the GCL (*p* = 0.13, **Fig. 3L**) or ML (*p* = 0.06, **Fig. 3L**) of the DG.

However, the C3 percentage area (**Fig. 3D-F, M,N**), representing the spread of neuroinflammatory processes, was more expansive, showing a significant main effect of genotype in both CA1 (F_(1, 36)_ = 23.8, *p* < 0.0001, **Fig. 3M**) and DG (F _(1, 8)_ = 9.6, *p* = 0.015, **Fig. 3N**). Post-hoc analysis demonstrated a widespread increase in C3 area coverage across the majority of CA1 laminae, including the SO (WT: 2.2 ± 0.42%, *n* = 6; PS19: 9.6 ± 3.5%, *n* = 5, *p* = 0.0078, **Fig. 3M**), SR (WT: 1.4 ± 0.3%, *n* = 6; PS19: 7.0 ± 2%, *n* = 5, *p* = 0.04, **Fig. 3M**), and SLM (WT: 1.7 ± 0.5%, *n* = 6; PS19: 10.3 ± 3.2%, *n* = 5, *p* = 0.002, **Fig. 3M**). Interestingly, the SP remained unaffected (*p* =0.13, **Fig. 3M**), suggesting that the cell body layer is relatively shielded from this neuroinflammatory transformation. In the DG, this expansion was localized to the GCL (WT: 1.7 ± 0.5%, *n* = 6; PS19: 10.3 ± 3.2%, *n* = 5, *p* = 0.002, **Fig. 3N**), leaving the ML unaffected (*p* = 0.21, **Fig. 3N**). Together, these data suggest that while GFAP-labelled astrogliosis is pan-hippocampal, the transition to a C3 neuroinflammatory state in the PS19 model is more laminar-specific.

### Microglial hyperactivity and laminar-specific changes in CD68

We next examined the microglial profile using antibodies to Iba1 (**Fig. 4A-C, G-J**) and the lysosomal/phagocytic marker CD68 (**Fig. 4D-F, K-N**). Two-way ANOVA revealed a significant main effect of genotype on Iba1 mean intensity in CA1 (F_(1, 16)_ = 31.8, *p* < 0.0001, **Fig. 4G**) and DG (F_(1, 8)_ = 13.32, *p* = 0.007, **Fig. 4H**) with PS19 mice exhibiting significantly higher expression compared to WT mice. Post-hoc analysis showed significant elevations in Iba1 intensity across all subfields, including the SO (WT: 51.8 ± 1.0, *n* = 6; PS19: 57.3 ± 2.3, *n* = 5, *p* = 0.03, **Fig. 4G**), SP (WT: 49.4 ± 1, *n* = 6; PS19: 57.7 ± 2.1, *n* = 5, *p* = 0.003, **Fig. 4G**), SR (WT: 52.4 ± 1.2, *n* = 6; PS19: 58.4 ± 3.0, *n* = 5, *p* = 0.02, **Fig. 4G**) and SLM (WT: 52.2 ± 1.0, *n* = 6; PS19: 59.0 ± 3.2%, *n* = 5, *p* = 0.01, **Fig. 4G**). Similarly, there were significant increases in both GCL (WT: 52.1 ± 1.9, *n* = 6; PS19: 59.1 ± 2.5, *n* = 5, *p* = 0.02, **Fig. 4H**) and ML (WT: 50.6 ± 1.0, *n* = 6; PS19: 56.5 ± 2.6, *n* = 5, *p* = 0.05, **Fig. 4H**) in the DG. However, in contrast to the GFAP findings, there were no significant differences in the percentage area of Iba1 immunolabeling across SO (*p* = 0.061, **Fig. 4I**), SP (*p* = 0.11, **Fig. 4I**), SR (*p* = 0.24, **Fig. 4I**), and SLM (*p* = 0.09, **Fig. 4I**) in CA1 subfields or in GCL (*p* = 0.2, **Fig. 4J**) or ML (*p* = 0.2, **Fig. 4J**) of DG.

**Figure 4.**
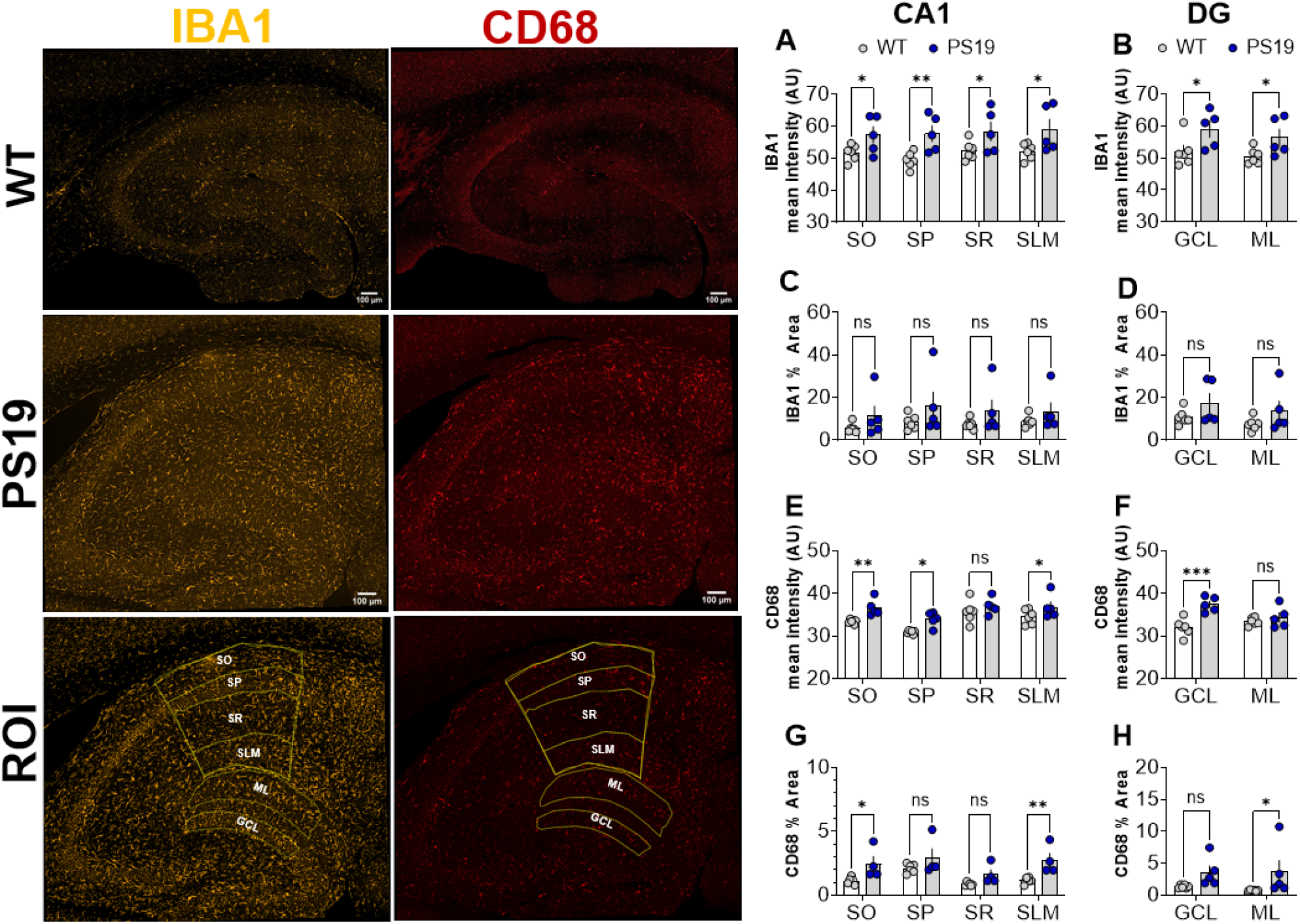
Widespread microglial activation and increased expression of the phagocytic marker CD68 in the PS19 hippocampus. **(A-C)** Representative confocal images of IBA1-positive microglia (yellow) in hippocampal sections from WT and PS19 mice. **(D-F)** Confocal images of CD68 expression (red), indicate lysosomal content within microglial compartments. **(C,F)** Regions of interest (ROI) for laminar quantification including CA1 subfields (stratum oriens, SO; stratum pyramidale, SP; stratum radiatum, SR; stratum lacunosum-moleculare, SLM) and dentate gyrus layers (molecular layer, ML; granule cell layer, GCL). **(G-J)** Quantitative analysis of IBA1 mean intensity and percent area. PS19 mice exhibited a significant increase in IBA1 intensity across all CA1 and DG layers, though the total area occupied by microglia remained relatively stable compared to WT. **(K-N)** Quantification of CD68 mean intensity and percentage area. PS19 mice showed significantly elevated CD68 expression, particularly in the CA1 SO and SLM, as well as the DG GCL, suggesting a shift toward a phagocytic microglial phenotype. Scale bars: 100 µm. Individual data points are an average of 3 sections per animal (n = 5-6 mice per group). All data presented as mean ± SEM; ns, p > 0.05; * *p* < 0.05; ** *p* < 0.01; *** *p* < 0.001.

For CD68 (**Fig. 4D-F, K-N**), there were significant genotype effects in mean intensities in CA1 (F_(1, 16)_ = 21.3, *p* = 0.0003, **Fig. 4K**) and DG (F_(1, 8)_ = 13.3, *p* = 0.007, **Fig. 4L**), with labelling intensity being higher in the PS19s compared to WT. Post-hoc analysis revealed that CD68 intensity was significantly elevated in the SO (WT: 33.4 ± 0.2, *n* = 6; PS19: 36.6 ± 0.9, *n* = 5, *p* = 0.008, **Fig. 4K**), SP (WT: 30.9 ± 0.2, *n* = 6; PS19: 33.9 ± 0.8, *n* = 5, *p* = 0.011, **Fig. 4K**), and SLM (WT: 34.6 ± 0.7, *n* = 6; PS19: 36.8 ± 1.2, *n* = 5, *p* = 0.05, **Fig. 4K**), but not in SR (*p* = 0.23, **Fig. 4K**). The DG GCL (WT: 32.0 ± 0.8, *n* = 6; PS19: 37.4 ± 0.8, *n* = 5, *p* = 0.0007, **Fig. 4L**) showed greater CD68 mean intensity but not the ML (*p* =0.36, **Fig. 4L**).

Two-way ANOVA indicated significant genotype differences in the percentage area of CD68 occupancy, both in CA1 (F_(1, 12)_ = 23.0, *p* = 0.0004, **Fig. 4M**) and DG (F_(1, 18)_ = 7.2, *p* = 0.02, **Fig. 4N**). Interestingly, the CD68 percentage area occupancy showed a divergent pattern between subfields: in the CA1, expansion was significant in the SO (WT: 1.1 ± 0.1, *n* = 6; PS19: 4.9 ± 2.5, *n* = 5, *p* = 0.01, **Fig. 4M**) and SLM (WT: 1.2 ± 0.1, *n* = 6; PS19: 4.6 ± 2.0, *n* = 5, *p* = 0.006, **Fig. 4M**), while no significant changes were observed in SP (*p* = 0.11, **Fig 4M**) or SR (*p* = 0.12, **Fig. 4M**). In the DG, the ML exhibited a marked increase in CD68 occupancy (WT: 0.7 ± 0.03, *n* = 6; PS19: 3.6 ± 1.8, *n* = 5, *p* = 0.05, **Fig. 4N**), but not the GCL (*p* = 0.115, **Fig. 4N**). The disassociation between the microglial markers suggests that although PS19 microglia exhibit increased activation and lysosomal density, they do not undergo the marked proliferation typically associated with proinflammatory/phagocytic states. Collectively, these data suggest general markers of heightened glial reactivity are widespread, while specific neuroinflammatory signaling (e.g., C3 intensity) is more anatomically localized to specific laminae.

## Discussion

The present study that 8–10-month-old PS19 mice exhibit an LTP deficit due to an aberrantly engaged metaplasticity and neuroinflammation caused by excessive tau accumulation. Our findings reveal that the heterodendritic and transregional metaplasticity mechanisms, which can serve to preserve the dynamic range of hippocampal neurons’ synaptic plasticity, are chronically occluded in the presence of advanced tauopathy. This functional failure is accompanied by a divergent physiological profile between the CA1 and DG and a laminar-specific neuroinflammatory signature, suggesting that tau-induced glial-neuronal interactions drive the circuit towards a ceiling effect of this form of metaplasticity.

Interestingly, our IO analysis in SR revealed an increase in basal transmission in the CA1 of PS19 mice, evidenced by increased fEPSP slopes. This observation is consistent with previous reports where hippocampal slices incubated with tau subsequently exhibited increased basal synaptic transmission (Brown et al., 2023; Wang et al., 2025). The absence of changes in the PPR in CA1 suggests the changes observed in basal transmission are postsynaptic in origin, potentially driven by tau-mediated mislocalization of glutamate receptors or alterations in dendritic excitability (Hoover et al., 2010). Previous studies have reported that GABAergic interneuronal function, which is critical for maintaining excitatory/inhibitory balance in the brain, is also impacted in tauopathies, whereby expression of mutant tau protein alone likely promotes the loss of hippocampal interneurons (Levenga et al., 2013; Loreth et al., 2012). However, this effect is unlikely to explain the IO changes we observed since a separate study showed that incubation with tau aggregates significantly enhanced basal synaptic transmission, but not paired-pulse facilitation, in the presence of GABA_A_ and NMDA receptor antagonists (Wang et al., 2025).

In contrast to CA1, the DG exhibited a markedly different profile. We observed reduced basal transmission at MML synapses, perhaps suggesting lower vesicle availability, fewer postsynaptic AMPA receptors, or fewer functional synapses. Given that there was a reduction in PPR selectively at the 20 ms and 30 ms inter-pulse intervals, a reduction in basal vesicle release probability is unlikely to explain the reduced basal transmission, which is normally associated with increased PPR. An alternative explanation for the reduced PPR at the early interpulse intervals could be an enhancement of feed-forward inhibition set up by the first pulse, given that GABAergic inhibition would not be completely blocked by the low dose of 0.2 μM gabazine employed in this study (Naylor & Wasterlain, 2005; Tuff et al., 1983). The degree of paired-pulse suppression from single and repetitive paired-pulse stimuli is highly dependent on the combination of stimulus intensity and interpulse interval; constant stimulus intensity with short interpulse intervals produces the greatest levels of paired-pulse suppression (Kapur et al., 1989; Waldbaum & Dudek, 2009). Another potential mechanism is the desensitisation of neurotransmitter receptors (Naylor & Wasterlain, 2005).

The impairment of LTP in the CA1 SR of PS19 mice observed here aligns with previous evidence that tau pathology disrupts LTP (Fá et al., 2016; Yoshiyama et al., 2007). However, our data on the early induction phase of LTP provides an added layer to this dysfunction. In the CA1 SR, PS19 mice showed a significant reduction in the first 5 minutes post-TBS, indicating that the very first stage of synaptic enhancement is severely compromised. Conversely, in the DG MML, the induction phase was non-significantly different between genotypes. This resilience of the early induction phase of LTP in the DG, despite a significant deficit in the later maintenance of LTP, highlights a fundamental difference in how these subregions handle tau-mediated toxicity. While CA1 exhibits a major breakdown of plasticity from the first seconds of stimulation, the DG appears to maintain its initial responsiveness, with deficits only emerging during the stabilization phase and even then, to a lesser extent than for CA1. Such resilience is further supported by evidence that, despite the early appearance and age-dependent accumulation of abnormal tau in the perforant path and DG granule cells, these circuits do not necessarily drive cognitive deficits until much later stages, such as 16 months (Harris et al., 2012).

Our investigation of metaplasticity effect provides a novel mechanistic explanation for the LTP impairments. In WT mice, priming stimulation in the SO recruits an astrocyte-dependent signaling pathway to heterosynaptically inhibit LTP in the SR and MML, a homeostatic brake that may be designed to prevent saturation of the synapses or to preserve recently learned information (Hulme et al., 2013). In PS19 mice, this priming effect was occluded; SO stimulation failed to induce further inhibition of LTP in CA1 SR or MML of DG. Crucially, we observed that SO LTP was also compromised, and in tandem with a robust neuroinflammatory profile marked by widespread astrogliosis and microglial activation. The elevation of the neurotoxic astrocyte marker C3 was most intense within the SO, which is the initiation site of the heterosynaptic metaplasticity, supporting the hypothesis that tau-induced glial activation mimics a chronic priming effect in CA1. We previously established that pharmacological TNF priming at high concentrations inhibits SO LTP (Singh et al., 2022), suggesting that the deficits in PS19 mice may result from gliosis-induced cytokine (e.g. TNF) release (Leyns & Holtzman, 2017; Singh et al., 2019). Notably, this tau-driven pathology mirrors certain aspects of the amyloid-beta (Aβ) mouse model, with both the APP/PS1 (Singh et al., 2019) and PS19 models showing impaired SO LTP in the priming pathway. However, the preservation of SO LTP observed during the acute application of Aβ (Hu et al., 2009; Zhao et al., 2018) may indicate that SO-specific mechanisms are more resilient to the immediate effects of exogenous Aβ than they are to the chronic neuroinflammation associated with prolonged Aβ and tau pathology. Further, the inhibition of LTP in the PS19 mice may be explained by the fact that C3 upregulation is significantly more extensive in TauP301S models (Wu et al., 2019), suggesting that C3-mediated neurotoxicity specifically drives the SO LTP deficits observed in 8–10-month-old PS19 mice. Lastly, the observed lack of significant difference in SO LTP during SO-MML-priming experiments likely stems from the experimental conditions rather than a true restoration of LTP in PS19 mice. Specifically, we previously demonstrated that MML-tetanization induces depotentiation of LTP in the SO priming pathway in mouse hippocampal slices with an intact CA3 circuitry (Sateesh et al., 2026).

The glial response observed in this study correlates with the recorded electrophysiological dysfunction in CA1 and DG, characterized by robust astrogliosis evidenced by increased GFAP area and intensity throughout all CA1 layers, consistent with a profound phenotypic shift in response to tau accumulation. Crucially, the spatial distribution of the complement component C3, a marker of neuroinflammatory A1 astrocytes (Liddelow et al., 2017), proved particularly revealing. Thus, while C3 area coverage increased broadly, its mean expression intensity was significantly elevated only within the SO. Given that SO is the locus of priming stimulation in our metaplasticity protocol, this localized increase in C3 mean intensity potentially mimics electrical priming activity to initiate neuro-glial communication and downstream signaling, thereby facilitating the subsequent inhibition of SR LTP. These results align with reports indicating that astrocytes are uniquely vulnerable to increased 4R tau isoforms in vivo and in vitro (Schoch et al., 2023). This results in dysfunction that may leave neurons prone to hyperexcitability and degeneration. Although 4R tau imbalance likely drives both neuronal and glial changes, tau-mediated toxicity within astrocytes may reveal unique mechanisms of disease progression (Schoch et al., 2023), suggesting that tau accumulation in the SO triggers a specific astrocytic gliotransmitter response that chronically activates heterosynaptic inhibitory signaling.

Our analysis of microglial markers in CA1 and DG revealed a specific dissociation between protein expression and morphological spread. We observed significant increases in the mean intensity of Iba1 and CD68 labelling, indicating a state of heightened activation and functional engagement. However, the percentage area covered by IBA1 remained unchanged, while a layer-specific increase in the percentage area of CD68 expression was observed within the SO, SLM, and ML. This suggests that while microglia in the PS19 hippocampus are functionally activated, likely processing pathological tau, they have not yet undergone the extensive proliferation or become reactive (microgliosis), often seen in advanced stages of neurodegeneration (Andersson et al., 2019; Luo et al., 2015; Yoshiyama et al., 2007), while CD68 is upregulated in regions of active phagocytosis. This phenotype may represent an intermediate inflammatory state that contributes to the priming of the astrocytic network without causing widespread structural disruption (Liddelow et al., 2017).

In conclusion, this study characterizes a specific functional deficit in the PS19 hippocampus, i.e., neuroglia-associated, region-specific impairments in synaptic plasticity. Further, by demonstrating that the metaplastic regulatory system is aberrantly engaged, we provide an additional framework for understanding why synapses fail to encode new information. The localized elevation of C3 in the SO and the constitutive engagement of heterosynaptic inhibition of LTP suggest an aberrantly engaged metaplasticity mechanism. Consequently, therapeutic strategies targeting the specific inflammatory mediators driving this metaplastic engagement, such as TNF, may hold promise for restoring hippocampus function in tauopathies such as FTD (Li et al., 2017; Ou et al., 2021).

## Acknowledgements

This research was funded by a grant from the Health Research Council of New Zealand (#22/177) to W.C.A and S.S.

## Author Contributions

S.S. and W.C.A. designed the research. W.C.A. and S.S. obtained funding to support the research. S.S. performed the *in vitro* experiments and analyzed the data. B.L. performed immunofluorescence experiments, confocal imaging, and analyzed the data. O.J provided tissue for immunofluorescence. S.S. wrote the manuscript, and S.S., B.L., O.J., and W.C.A reviewed and amended the manuscript and approved the final submission.

