## Supporting information for "Synaptic plasticity deficits via aberrant engagement of metaplasticity in the hippocampus of PS19 mice"

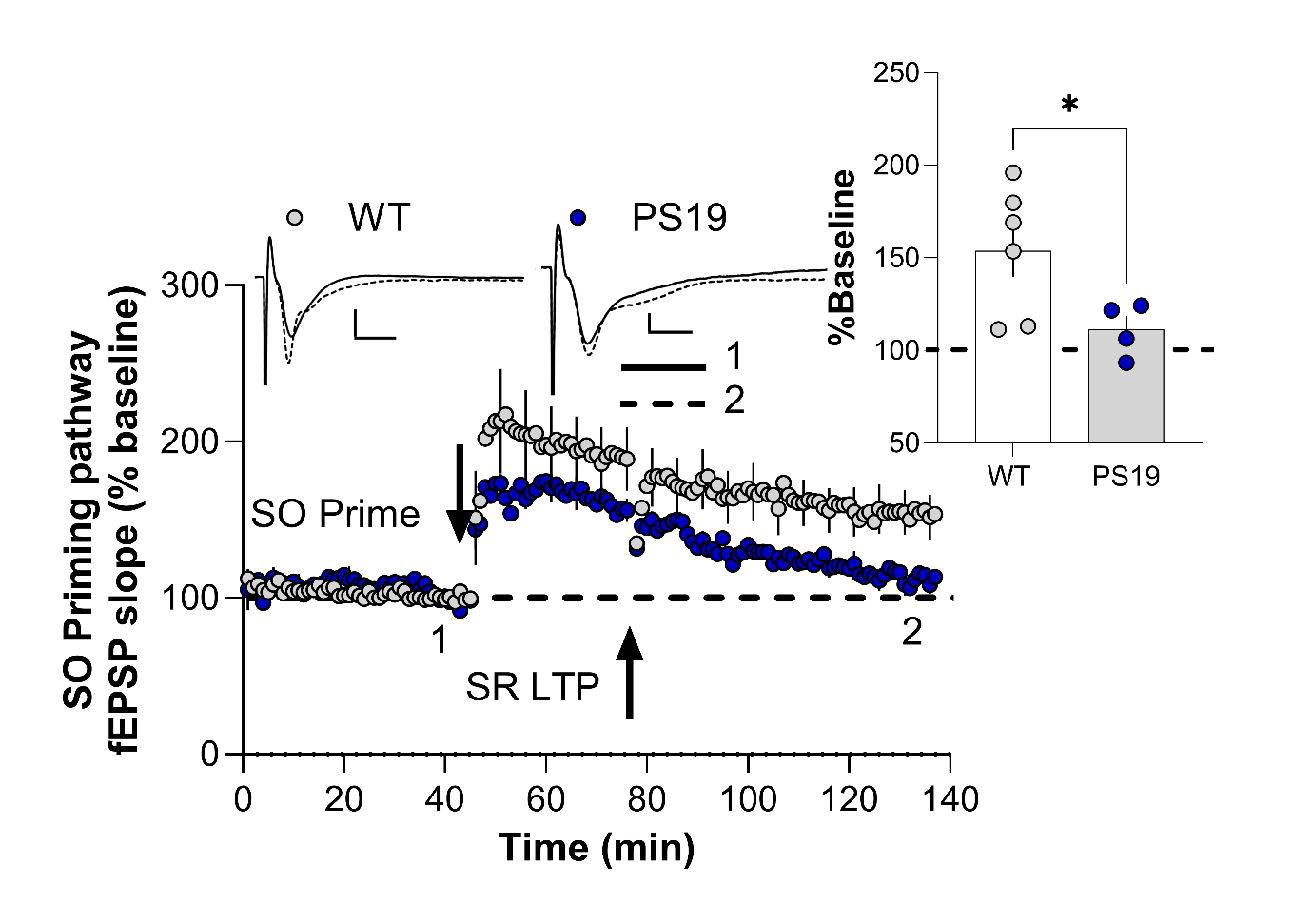

**Supplementary Figure 1: Impaired LTP in the SO priming pathway of PS19 mice.**

SO priming resulted in substantial homosynaptic LTP in WT mice; however, this potentiation was significantly attenuated in PS19 littermates. *Insets:* Representative fEPSP waveforms recorded prior to induction (**1**) and at the conclusion of the experiment (**2**). Arrows indicate the timing of SO priming and SR LTP induction. Scale bar: (WT) 0.5 mV, 5 ms ; (PS19) 0.2 mV, 5 ms. All data presented as mean ± SEM; * *p* < 0.05.

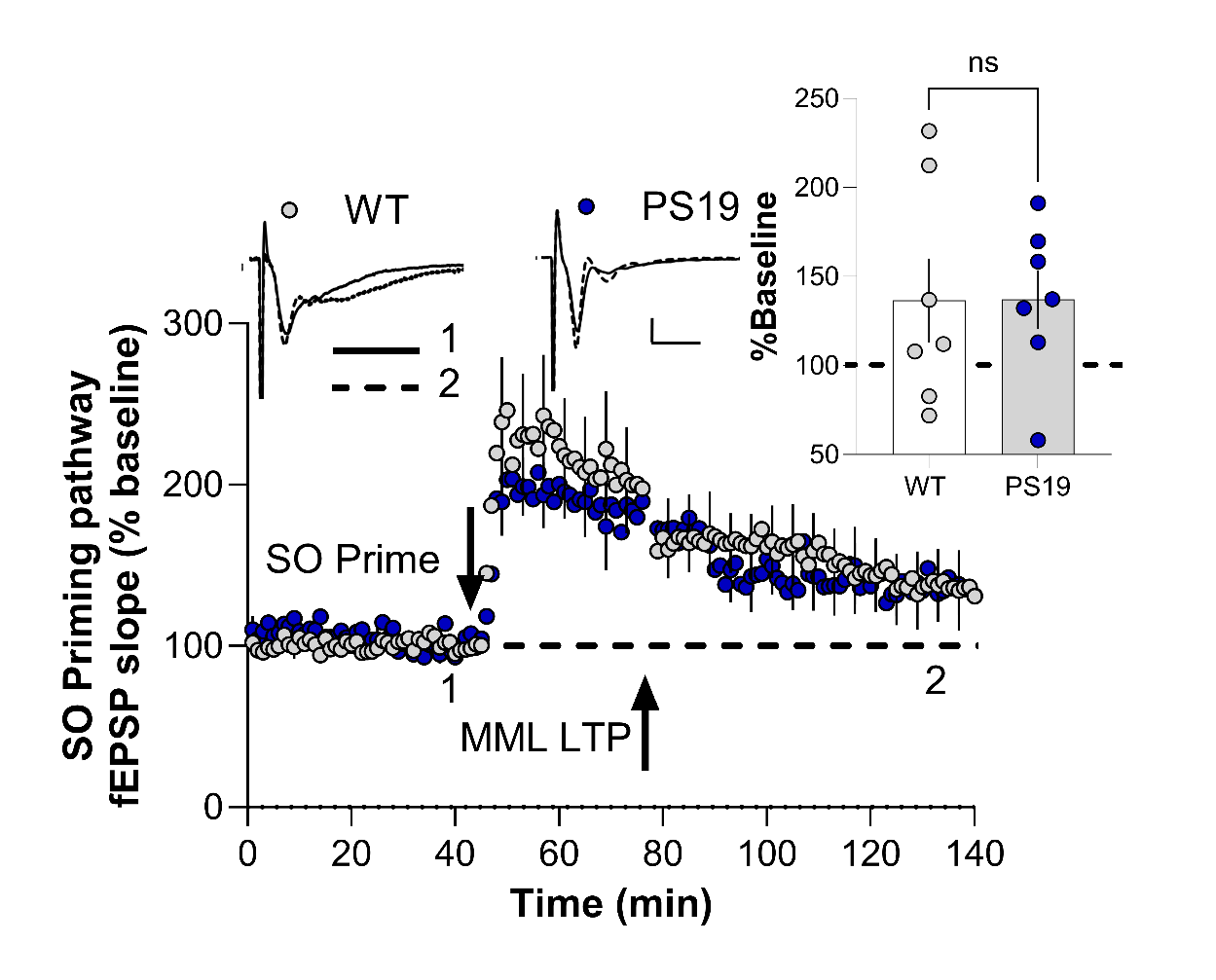

**Supplementary Figure 2: No significant difference in SO priming pathway LTP between PS19 and WT mice in the presence of 0.2 µM gabazine**.

SO priming resulted in substantial homosynaptic potentiation in WT and PS19 mice. Both genotypes exhibited equivalent levels of LTP stabilization at the conclusion of the recording period. *Insets:* Representative fEPSP waveforms recorded prior to induction (**1**) and at the conclusion of the experiment (**2**). Arrows indicate the timing of SO priming and MML LTP induction. Scale bar: 0.5 mV, 5 ms. All data presented as mean ± SEM; ns: *p* > 0.05.
